# A modified FASP protocol for high-throughput preparation of protein samples for mass spectrometry

**DOI:** 10.1101/084533

**Authors:** Jeremy Potriquet, Marut Laohaviroj, Jeffery Bethony, Jason Mulvenna

## Abstract

To facilitate high-throughput proteomic analyses we have developed a modified FASP protocol which improves the rate at which protein samples can be processed prior to mass spectrometry. Adapting the original FASP protocol to a 96-well format necessitates extended spin times for buffer exchange due to the low centrifugation speeds tolerated by these devices. However, by using 96-well plates with a more robust polyethersulfone molecular weight cutoff membrane, instead of the cellulose membranes typically used in these devices, we could use isopropanol as a wetting agent, decreasing spin times required for buffer exchange from an hour to 30 minutes. In a typical work flow used in our laboratory this equates to a reduction of 3 hours per plate. To test whether our modified protocol produced similar results to FASP and other FASP-like protocols we compared the performance of our modified protocol to the original FASP and the more recently described eFASP and MStern-blot. We show that all FASP-like methods, including our modified protocol, display similar performance in terms of proteins identified and reproducibility. Our results show that our modified FASP protocol is an efficient method for the high-throughput processing of protein samples for mass spectral analysis.

**Abbreviations**
ABC: Ammonium bicarbonate
ACN: Acetonitrile
TFA: Trifluoro acetic acid
FA: Formic acid
TEAB: Triethylammonium bicarbonate
DCA: Deoxycholic acid
DTT: Dithiothreitol
IAA: Iodoacetamide

## Introduction

Filter-aided sample preparation (FASP) is a method for efficiently generating tryptic peptides from complex protein mixtures prior to mass spectral analysis [1]. FASP uses a molecular weight cut-off (MWCO) membrane as a ‘reactor’ on which complex protein mixtures can be chemically modified and digested. The filter unit allows, in the presence of concentrated urea, the efficient removal of detergents and excess of reagents involved in chemical modifications of the proteins mixtures [2]. In FASP, microlitre-volumes of protein mixtures are retained on ultrafiltration devices while detergents and other low molecular weight impurities are depleted in buffers containing concentrated urea before chemical modification of thiols, protein digestion and peptide elution from the filtration device [1]. FASP therefore provides an efficient, convenient and effective method for the processing of cell or tissue lysates containing detergents [3] and has become widely adopted part of proteomic work flows [4, 5].

Although a clever solution to the preparation of protein samples for mass spectrometry, the original FASP methodology used individual ultrafiltration devices, such as Microcon (Millipore) or Vivacon (Sartorius-Stedim), that are resistant to reasonably high centrifugation speeds. However, for high-throughput processing of many protein samples, the use of cellulose MWCO filters, as used in the FASP protocols, in a 96-well plate format is problematic due to the low centrifugation speeds tolerated by these devices. Centrifugation between wash steps then becomes prohibitively long, drastically increasing the time needed for the processing of samples [6,7]. To facilitate high-throughput analysis of clinical samples, we have developed a method that uses a FASP-like protocol on 96-well plates using a more robust polyethersulfone (PES) filtration membrane in place of the cellulose filter typically used in FASP protocols. The use of this membrane, in conjunction with isopropyl alcohol as a wetting agent provides a 96-well FASP method which greatly reduces the time required for high-throughput processing of protein samples. The use of PES also provides greater flexibility in the choice of organic solvents during sample preparation and a reduced risk of sample loss after perforation [8] due to the greater chemical and physical robustness of the PES membrane.

Since the original description of FASP in 2009 [8], a number of modified protocols have been proposed to improve performance: enhanced FASP (eFASP) pre-passivates Microcon filter surfaces with 5% TWEEN-20 to enhance peptide recovery and uses a surfactant (0.2% deoxycholic) during detergent steps and digestion to increase trypsin efficiency [9]; 96-well plates with a 10 MWCO membrane have been used to enable economic high-throughput processing of urine and human cell lines [6, 7]; and, more recently, MStern-blot (MStern) uses a large-pore, hydrophobic polyvinylidene fluoride (PVDF) membranes in a 96-well format for improved high-throughput peptide recovery [10]. To evaluate the relative performance of our protocol compared to the other FASP-like methods and to assess any potential effects on proteins identified in subsequent mass spectrometry, we compared the performance of our modified protocol with the other FASP-like methods, FASP, eFASP and MStern-blot in the proteomic analysis of a complex protein mixture from human cell lines.

## Results and Discussion

### Polyethersulfone as a membrane for FASP analysis

Processing of tissue samples in 96-well plates is a convenient and high-throughput method for processing large numbers of samples prior to mass spectrometry, particularly for clinical applications. However, using FASP in 96-well plates equipped with cellulose MWCO membranes typically requires centrifugation spin times of an hour or more at 3200 × g for efficient buffer exchange in wash steps [6, 7, 10]. We modified the standard FASP protocol to use more robust PES filtration membrane and were then able to introduce 10% isopropanol into our work flow as a wetting agent (Suppl. Fig. 1), resulting in a reduction of ~50% in the time required to achieve full buffer exchange at 3,200 × g. With at least seven buffer exchange steps in our work flow (Suppl. Fig. 1) this amounted to a reduction of 3 hours in sample preparation time. The ability of isopropanol to improve the rate of buffer exchange is likely due to a reduction in surface tension between the aqueous layer and the membrane [11] and although alcohol can reduce the critical micelle concentrations (CMC) of detergents [12], affecting both the elimination of the detergent during buffer exchanges and its ability to solubilize proteins, this is likely balanced by the positive effects of urea on the CMC [13]. Using this modified protocol we have now successfully processed protein samples from a wide variety of sources including cell lines, bile extracts, human tissue and exosome preparations.

### Proteins identified using different processing methods

PES 96-well plates are available with both 10 and 30 kDa MWCO membranes and to compare the performance of our modified protocol to the other FASP-like protocols, we used each method, FASP, eFASP, MStern and the 30 and 10 kDA versions of our method (referred to below as pFASP-30 and −10), to process total protein from a cancer cell line (KKU055) before tandem mass spectral analysis on an AB SCIEX 5600+ mass spectrometer. Figure 1 summarizes the number and characteristics of proteins identified using the different methods. The modified protocol provided a comparable number of protein identifications as the other FASP-like methods; identified proteins ranged from 741 for eFASP to 644 for MStern (Figure 1A) with pFASP-10 and −30 providing 675 and 722, respectively. Each method showed similar reproducibility as measured by the number of unique protein identifications in each replicate; MStern providing the most reproducible set of protein identifications (11%) and pFASP-10 the least (14%; Figure 1B).

**Figure 1:**
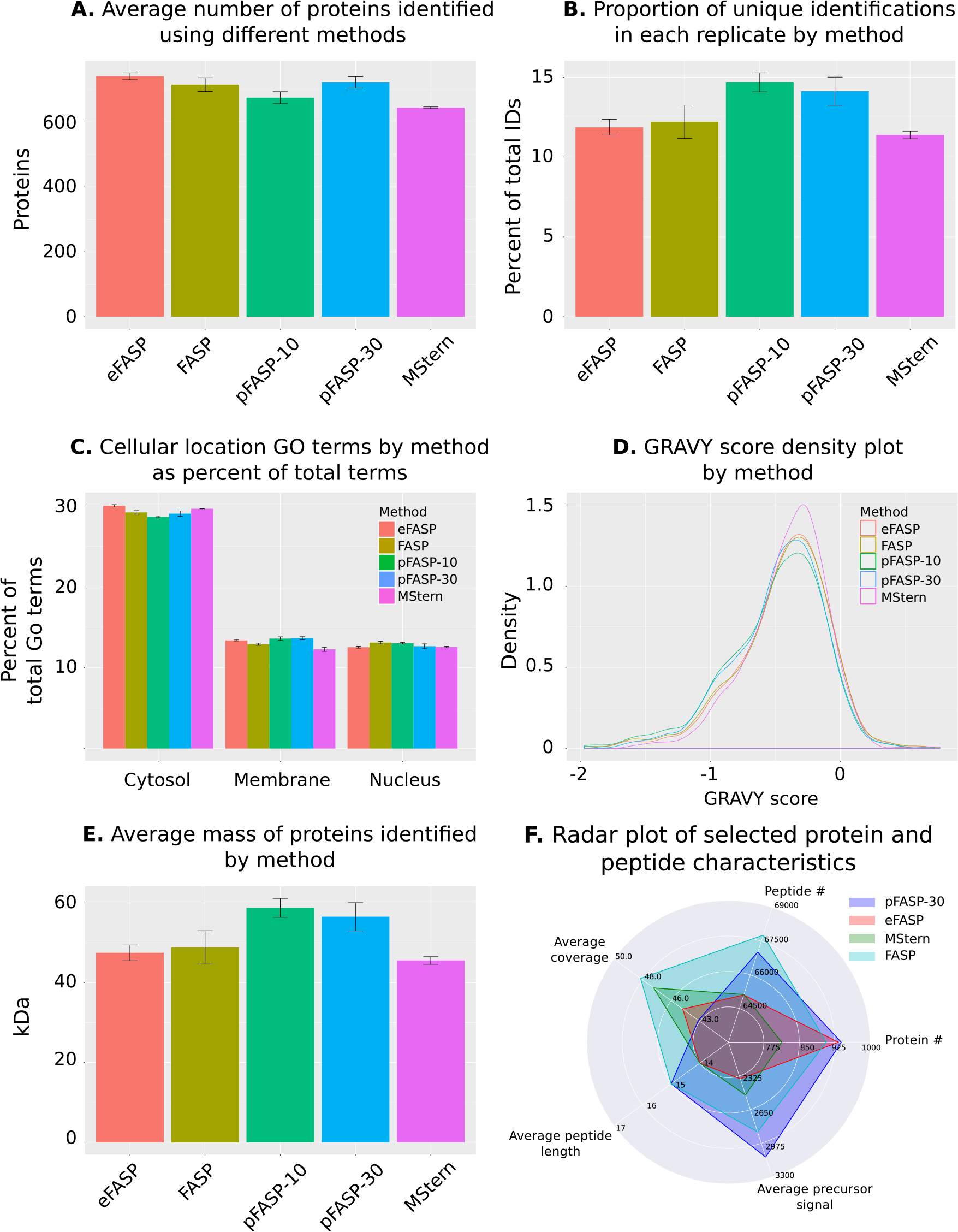
Total protein identification using different processing methods. **A.** Bar plot comparison of the average number of non-redundant proteins identified across three replicates of each sample preparation method; **B.** Bar plot comparison of the average number of unique proteins identified in each replicate of each method; **C.** Average percent of total ‘cellular component’ GO terms returned by proteins identified in each replicate of each method that were associated with a cytosolic, membrane or nuclear subcellular location; **D.** Density plots of GRAVY scores for proteins identified using each protein processing method; **E.** The average mass of proteins identified in each replicate of each protein processing method; and **F.** Radar plot showing selected characteristics of proteins and peptides identified using each protein processing_12_ method.

### Cellular location and hydrophobicity of identified proteins

To determine if there were differences in the characteristics of proteins identified using different FASP methods the cellular location, hydrophobicity and size of identified proteins were compared. A comparison of the ‘cellular component’ GO terms associated with identified proteins suggests that the pFASP methods identified slightly more membrane proteins than the other methods (Figure 1C) although differences between each method were minor. Likewise, density plots of protein GRAVY scores (a measure of the hydropathy of amino acid sequences [14]) were quite similar although MStern did show fewer protein identifications in the hydrophilic regions of the plot (Figure 1D). As the only method that relies on hydrophobic interactions rather than a MWCO to retain proteins on the membrane, this could reflect a tendency for more hydrophilic proteins to pass through the PVDF membrane before the completion of processing, as previously suspected [10] and as reflected in the lower number of membrane proteins identified using this method (Figure 1C).

### Unique identifications provided by different methods

To determine whether each method identified a similar set of proteins, protein identifications from all replicates were combined and the overlap in protein identifications visualized using an Upset plot [15]. From 1,669 identified proteins, only 403 were identified in all methods (Figure 2A). Some 542 protein identifications, 32% of the total, were identified in only one of the tested methods (highlighted in Figure 2A). Although, in this analysis, without pre-fractionation of proteins and relatively short chromatographic separation prior to MS analysis, this could be explained by stochastic effects, the GRAVY scores and cellular location GO terms for each method were compared to determine if there were any systematic effects associated with different methods. GRAVY score density plots for identified proteins in each method suggests that the MStern method provided more unique protein identifications within a narrower range of hydrophobicity, with the other methods providing unique identifications of both more hydrophilic and hydrophobic proteins. As discussed, more hydrophilic proteins may not be retained on the PVDF membrane but it is also possible that some hydrophobic peptides may not elute from the membrane after processing, as evidenced by the longer right-hand tails of the MWCO methods in the GRAVY score density plot (Figure 2B). In combination with the GRAVY score density plot in Figure 1D, it appears that MStern is providing deeper coverage over a smaller range of hydrophobicity while the MWCO methods provide identifications over a greater range of hydrophobicity at the cost of a shallower coverage (under these experimental conditions). This was also reflected in both the greater number of membrane-associated GO terms provided by the unique identifications from the MWCO methods (Figure 2C) and the further 136 protein identifications not identified using the MStern method (marked with asterisks in Figure 2A).

**Figure 2:**
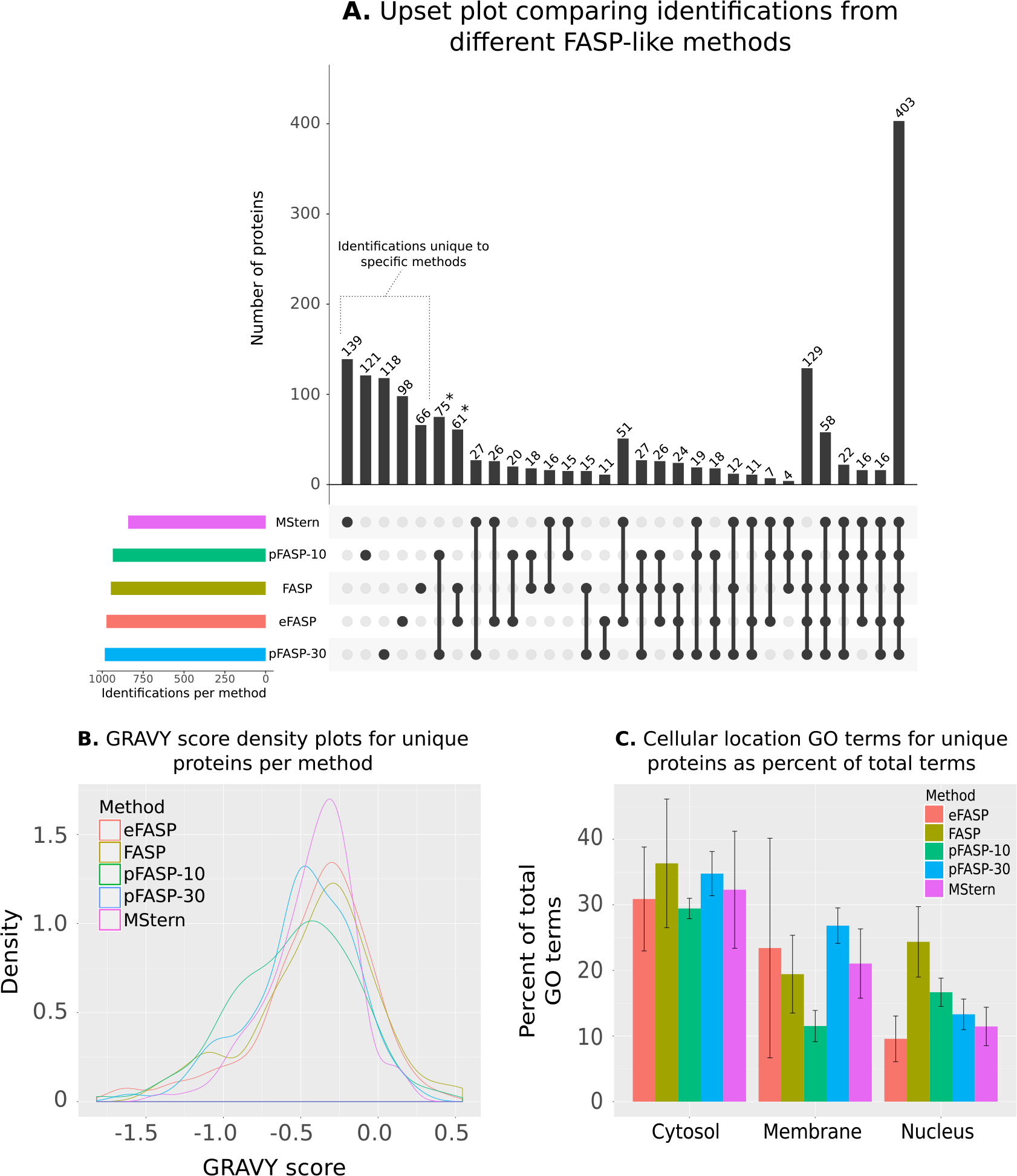
Upset analysis of proteins identified using each sample preparation method and characteristics of unique proteins only identified using a single method. **A.** Upset chart showing overlap in proteins identified using each protein processing method. Numbers of identified proteins shared between different sets of methods are indicated in the top bar chart and the specific methods in each set are indicated with solid points below the bar chart. Total identifications for each method are indicated on the left as ‘Identifications per method’. Figure generated using Upset R package [15]; **B.** Density plots of GRAVY scores for proteins uniquely identified in each protein processing method; and **C.** Average percent of total ‘cellular component’ GO terms returned by proteins uniquely identified in each replicate of each method that were associated with a cytosolic, membrane or nuclear subcellular location.

### Conclusions

In this work we have shown that our modified FASP protocol is a useful adaptation of the FASP protocol for decreasing the time need for the high-throughput analysis of large numbers of samples in a 96-well format. Using our method we were able to decrease spin times by 50%, representing 3 hours in our typical sample preparation work flow. When compared to the different FASP-like sample preparation protocols our method provides similar performance to other FASP protocols with a similar number of protein identifications and the same level of reproducibility. In our modified protocol we exploit the increased robustness of the PES membranes to use isopropanol as a wetting agent allowing for faster buffer exchange, but the tolerance of the membrane for a range of organic solvents allows for other potential work flows using these solvents, for example the processing of proteins in phenol solution after DNA extraction. We have now used our modified method for the processing of protein mixtures from a wide-range of biological sources and the work presented here suggests our method provides a useful technique for high-throughput processing of samples for proteomic analysis.

## Experimental

The same lysis and protein solubilisation protocol was observed for all methods. Removal of reduction and alkylation agents were performed on the filter devices as described in the respective protocols for each technique. All tubes and solvent containers used were washed with 50% methanol and dried to minimize potential polyethylene glycol contamination.

### Lysis and protein solubilisation of H69 normal human cholangiocyte cells

Adherent H69 Human cells were harvested using a 5 min incubation with TrypLE express solution (Gibco) at 37°C followed by three washes with Dulbecco’s PBS (Gibco) with centrifugation at 800 × g for 5 min between each wash. Approximately 6 million cells were lysed on ice by resuspending a cell pellet in 250 *μ*L of 1% SDS, 5 mM MgCl_2_, 10 mM CHAPS and 100 mM TEAB supplemented with 1 × Roche Complete protease inhibitors. Immediately after lysis, DNA and RNA were degraded by the addition of 50 mM Tris, 20 mM NaCl and 2 mM MgCl_2_ with 10 *μ*l of 1 unit/*μ*l ultrapure Benzonase (Sigma) followed by incubation at 4°C for 30 minutes with constant agitation. Protein concentration was determined using the BCA Assay (Pierce) following the manufacturer’s protocol with a standard curve of 1 mg/ml, 0.5 mg/ml, 0.25 mg/ml, 0.125 mg/ml and 0.0625 mg/ml from a 2 mg/ml BSA stock solution. After 30 min of incubation at 37°C, the measurement was performed in a microplate spectrophotometer (Synergy Hybrid H4 from Millennium Science) at a wavelength of 562 nm. Aliquots of 50 × g of proteins in 10 *μ*L lysis buffer were transferred into individual 1.7 mL microtubes (Axygen) in sufficient number to perform triplicates of each sample processing technique.

### Modified FASP protocol (10 and 30 kDa)

Proteins in lysis buffer were reduced by the addition of 0.5M DTT stock solution to a final concentration of 20 mM followed by incubation at 95°C for 5 min and cooling at rt for 10 min. Proteins were then alkylated in 40 mM of IAA for 45 min in darkness. Eight volumes of 8M Urea, 10% isopropanol in 100 mM TEAB was then added to the protein sample. Either 10 or 30 kDa molecular weight cut-off filter plates (AcroPrep advance 96-well Omega filter plates, PALL), coupled to deep U-bottom well plates (Axygen) for collection, were prepared by briefly spinning 200 *μ*L of 60% isopropanol through the filter at 3,100 × g. Protein sample was then transferred to the plate and spun at 3,100 × g for 30 min in a 5810R (Eppendorf) centrifuge with adapted plate bucket. Detergent removal by buffer exchange was performed in two successive washes with 8M urea, 10% isopropanol in 100 mM TEAB with centrifugation at 3,100 × g for 30 min between each wash. Urea was then removed by two washes with 10% isopropanol in 50 mM TEAB with centrifugation at 3,100 × g for 30 min between each wash. A final of wash with 50 mM TEAB was performed with centrifugation as described above. Protein digestion was then performed by adding 1 × g of trypsin, in 50 mM TEAB, to the wells and incubating overnight at 37°C. Peptides were recovered using an initial spin of 3,100 × g for 10 min followed by two centrifugations with 50 *μ*L of 50 mM TEAB. Recovered peptides were dried in a speed vacuum for 4 hours at 45°C.

### FASP

FASP was performed as previously described [1] with minor changes. Briefly, proteins were reduced and alkylated as described for above and were then mixed with eight volumes of 8M urea in 100 mM Tris-HCl pH8. Microcon Ultracel 30 kDa (Millipore) units were prepared by briefly spinning 60% methanol through the filter at 14,000 × g. Protein samples were transferred to the filter units and spun at 14,000 × g for 15 min. Detergent removal by buffer exchange was performed in two successive wash with 8M urea in 100 mM Tris-HCl pH8 with a 15 min spin at 14,000 x g. Unlike the original protocol, three additional washes using 100 mM Tris-HCl pH8, with a 15 min spin at 14,000 × g between each wash, were included to remove excess urea. Protein digestion was achieved by adding 1 × g of trypsin in 50 mM ABC and incubating at 37°C overnight. Peptides were recovered in two washes with 50 *μ*L of 50 mM ABC with spinning at 14,000 × g for 5 min each. The mixture was then vacuumed dried for 4 hours at 45°C.

### eFASP

eFASP was performed as previously described [9] with minor changes. Briefly, the day before the experiment a Microcon Ultracel 30 kDa filtration unit (Millipore) was “passivated” by an overnight incubation in a solution of ultrapure water with 5% v/v of TWEEN-20. Proteins in lysis buffer were then reduced and alkylated as described above and mixed with eight volumes of 8M urea, 0.2% DCA in 100 mM Tris-HCl pH8. The passivated filter units were rinsed thoroughly by three immersions in a large volume of ultrapure water and the protein mixtures transferred to them before spinning at 14,000 × g for 15 min. Detergent removal by buffer exchange was done using two successive washes of 8M urea, 0.2% DCA in 100 mM Tris-HCl pH8 with spinning at 14,000 × g for 15 min after each wash. Urea was then removed in three washes of 50 mM ABC with 0.2% DCA with spinning at 14,000 × g for 15 min. Protein digestion was achieved by adding 1 × g of trypsin in 50 mM ABC with 0.2% DCA and incubating at 37°C overnight. Peptides were recovered in two washes with 50 μL of 50 mM ABC with spinning at 14,000 × g for 5 min. Peptides were then lyophilised prior to mass spectral analysis.

### MStern-Blot

MSTern-Blot was performed as previously described [10] with minor changes. Briefly, proteins were reduced and alkylated as described above and mixed with eight volumes of 8M urea in 100 mM ABC. The wells of a 0.45 μm hydrophob-high protein binding immobilon-P membrane (PVDF) plate (Millipore) were prepared by passing 70% ethanol through the filter on a vacuum manifold on top of a deep U-bottom well collecting plate. The PVDF membrane was equilibrated with 8M urea in 100 mM ABC solution before application of the protein mixture to the filter plate wells. The protein mixture was passed through the membrane three times by collecting the flow through and re-applying it to the filter membrane. Unlike the original protocol, an extra wash using 8M urea in 50 mM ABC was used prior to two washes with 50 mM ABC. Protein digestion was performed by adding 1 × g of trypsin in 50 mM ABC and incubating at 37°C overnight. Peptides were recovered in a V-bottom well collecting plate using two washes with 40% ACN, 0.1% TFA in MS-grade water and lyophilised prior to mass spectral analysis.

### Tandem mass spectrometry

Tryptic peptides were solubilised in a solution of 0.1% TFA and desalted on ZipTip C18 pipette tips (Millipore). Desalted peptides were lyophilised and resuspended in 30 μL of 0.1% formic acid [aq], 2% ACN and centrifuged at 12,000 × g for 5 min to remove particulate matter. Peptides were chromatographically separated on a Eksigent Ekspert nanoLC 415 (AB SCIEX). Firstly, 2 l of desalted peptides were injected onto a ChromXP C18CL trap column (3 μm, 120 Å, 10 x 0.3 mm) at 5 l/min for five min in 0.1% formic acid [aq]/5% ACN (solvent A) before being placed in line with a ChromXP C18 (3 μm, 120 Å, 15 cm x 75 μm) nanoLC column. Peptides were elute by three consecutive linear gradients: 5-10% solvent B (ACN/0.1% formic acid) over 2 min, 10-40% solvent B over 58 min and 40-50% solvent B over 5 min at a 300 nL/min flow rate. Eluted peptides were passed directly into a nano-electrospray ion source of a TripleTOF 5600+ mass spectrometer (AB SCIEX) for tandem mass spectrometry with the following parameters: ion spray voltage was set to 2300V, declustering potential of 150 V, curtain gas flow 25, nebuliser gas 1 (GS1) 15 and interface heater at 150°C. Full scan TOF-MS data was acquired in Information Dependant Acquisition (IDA) mode over the mass range 350-1350 (m/z) with 250 ms accumulation time and, for product ions, 100-2000 (m/z) with 50 ms accumulation time for a total of 10,606 cycles. Ions observed in the TOF-MS scan, exceeding a threshold of 150 counts and possessing a charge state of +2–+5, were set to trigger the acquisition of product ion spectra for a maximum of 40 of the most intense ions. Dynamic exclusion was incorporated for 10 seconds after one occurrence of a precursor ion. The mass spectrometer was automatically recalibrated with B-Galactosidase digest after every 2 samples. Data was acquired and processed using Analyst TF 1.7 software (AB SCIEX).

### Database searching

Spectral searches of processed LC-MS/MS data were performed using ProteinPilot v4.5 (AB SCIEX) using the Paragon algorithm (version 4.5.0.0) with the following parameters: one miss cleavage allowed, carboxymethylation was specified as a fixed modification and oxidation of methione as a variable modification, urea denaturation was also set as special factors. Background correction was used and biological modifications specified as an ID focus. The detected protein threshold was set as 0.5 and the false-discovery rate (FDR) was calculated using searches against a decoy database comprised of reversed sequences. Searches were conducted against the SwissProt human reference proteome set comprising 70236 protein sequences (downloaded 30th, March 2016).

## Acknowledgments

This research was supported by award R01CA155297 from the National Cancer Institute, award P50AI098639 from the National Institute of Allergy and Infectious Disease and fellowship support (JPM) and research support (grant number 1051627) from the National Health and Medical Research Council of Australia. JM is supported by a Career Development Fellowship from the National Health and Medical Research Council, Australia (NHMRC).

